# SPOUT: An open-source hardware and software platform to study decision making while manipulating and recording from neural activity

**DOI:** 10.64898/2026.08.21.746318

**Authors:** Bastijn J.G. van den Boom, Deeptirmayee Dash, Melanie Rutherford, Allison E. Girasole, Pavel Gorelik, Ofer Mazor, Bernardo L. Sabatini

## Abstract

Recording and manipulating brain activity during behavior is critical to understanding the underlying mechanisms of decision-making. Linking neural activity to behavior requires behavioral hardware and software tightly integrated with recording and perturbation systems on a shared clock. We built SPOUT (State-machine Platform for Operant Uni/dual-spout Tasks), an open-source, Teensy-driven state-machine platform with a MATLAB interface that runs 10 unique decision-making tasks (with dozens of variations available through user-friendly settings) to study behavior in head-restrained mice. The platform is built on several custom hardware devices: a dual-lick detector, headplate designs for optogenetics and two-photon calcium imaging, a three-axis motorized spout manipulator, and an optogenetics power modulator. The firmware differentiates between one and two lick spout tasks and can be controlled by a user-friendly interface. Task settings can be selected through the interface or by loading predefined settings files. We validated the clock speed and lick detection against an independent, external acquisition system and identified highly precise, sub-millisecond detection of single licks. Using a pseudo-random synchronization pulse generated by SPOUT, we corrected for missing data due to glitches in the acquisition system and clock drift. We showcase the versatility of SPOUT by training mice on an uninstructed lick-left/lick-right task in which the rewarded side switches unexpectedly and found that mice use history-dependent action-outcome associations to guide future behavior. Transiently inhibiting the anterior lateral motor cortex (ALM) during cue presentation induced contralateral deficits, without affecting ipsilateral trials. Finally, two-photon imaging of ALM neurons revealed stronger population responses during contralateral choice licks compared to ipsilateral ones. Together, SPOUT offers an open-source, affordable platform to study decision-making in head-restrained mice while combining neural recordings and manipulations.

## INTRODUCTION

Understanding how neural activity gives rise to behavior requires experimental approaches that combine precisely quantified behavior with simultaneous recording and manipulation of brain activity. Head-restrained behavioral paradigms are central to this effort because they provide the stability required for modern optical and electrophysiological techniques while maintaining the experimental control needed to measure behavior on a trial-by-trial basis^1–6^. These preparations permit large-scale recordings with two-photon calcium imaging^7–9^ and high-density electrophysiology^6,10,11^, and enable causal interrogation of neural circuits using optogenetics^12–14^. As behavioral paradigms and neuroscience technologies become increasingly sophisticated, state machines need to implement higher temporal precision to resolve single events and be able to synchronize behavior to neural activity with high accuracy^15^.

Building such precise and accurate behavioral monitoring and control systems remains expensive and technically challenging. Behavioral hardware must detect animals’ actions with sub-millisecond latency, execute task logic deterministically, and record all behavioral events. At the same time, behavioral timestamps must be aligned with external acquisition systems that often operate on independent clocks^16^. Even small discrepancies arising from dropped samples, acquisition glitches, or gradual clock drift can complicate the alignment of behavioral and neural data, particularly during long recording sessions.

Furthermore, many laboratories develop custom hardware and software tailored to individual experiments, making it difficult to reproduce behavioral paradigms across setups and laboratories or to reuse task logic across different studies. Although several open-source behavioral platforms have substantially lowered the barriers to implementing behavioral experiments^2–4,17–19^, they often emphasize specific behavioral paradigms, recording modalities, or hardware configurations, requiring additional customization to combine multiple decision-making tasks with closed-loop neural perturbation and recording. Increasing attention to reproducibility across behavioral neuroscience further highlights the need for behavioral platforms that are modular, transparent, and easily shared across laboratories^20,21^.

To address these challenges, we developed SPOUT (State-machine Platform for Operant Uni/dual-spout Tasks), an open-source hardware and software platform designed for studying decision-making in head-restrained mice while integrating seamlessly with behavioral cameras, neural recordings and perturbation techniques. SPOUT combines Teensy microcontroller-based firmware architecture with modular hardware and a user-friendly MATLAB graphical interface to provide flexible control over behavioral experiments without requiring changes to the underlying firmware or programming experience. The Teensy state machines, in essence models that navigate through a fixed number of conditions by means of transitions in response to external events, allow for sub-millisecond detection of behavioral events. The platform supports 10 commonly used behavioral tasks using two generalized state machines for one- and two-spout paradigms, with dozens of additional task variants available through configurable settings rather than new firmware or code. SPOUT provides temporal precise, closed-loop optogenetics stimulation through any external light source. Hardware developments include an electrically isolated dual-spout lick detector, motorized spout positioning, headplate designs compatible with optogenetics and high aperture objective two-photon imaging, and a calibrated closed-loop optogenetic stimulation module. To facilitate integration with external acquisition systems, SPOUT generates pseudo-random synchronization pulses that allow robust alignment of behavioral events with independently acquired neural data and behavioral camera frames, while remaining resilient to missing samples and clock drift.

Here, we describe the design and validation of the SPOUT platform. We first characterize the hardware, software architecture, timing performance, lick detection, and synchronization strategy. We then demonstrate the versatility of the platform by training mice on a history-dependent lick-left/lick-right decision-making task, demonstrating that animals acquire action-outcome associations that guide future choices^22–25^. Next, we use integrated closed-loop optogenetics stimulation to transiently inhibit anterior lateral motor cortex (ALM) during task performance, reproducing the well-established contralateral behavioral deficits^1,7^. Finally, we combine SPOUT with two-photon calcium imaging to relate neuronal population activity in ALM to behavioral choice. Together, these experiments establish SPOUT as an affordable, user friendly, open-source platform for implementing decision-making experiments while enabling precise manipulation and recording of neural activity.

## RESULTS

### Hardware for head-restrained decision-making tasks in mice

Behavioral neuroscience experiments that combine perturbation or recording of neural activity with detailed behavior measurements depend on hardware and software that respond to an animal’s actions with sub-milliseconds latency, log every event, and produce a synchronization signal that allows matching of different clocks. To achieve this precision, we developed an open-source platform, the State-machine Platform for Operant Uni/dual-spout Tasks (SPOUT), from custom designed and off-the-shelf components chosen for ease of reproduction (Figure 1A). A single setup costs ∼ $1,000 in parts (Supplementary figure 1), excluding a computer and screen. A breadboard carries connections for power supplies (5, 3.3, and 12 V) that power a custom designed Teensy breakout board with integrated solenoid drivers (5 V, Supplementary figure 2), a conduction-based dual-lick detector (3.3 V, Supplementary figure 3), and an audio amplifier (12 V) and speakers for auditory cues, respectively. Water is delivered through solenoids fed by reservoir syringes and cleared through flush syringes via 3-way valves. The dual-lick detector follows a conduction-based design in which contact with a grounded spout pulls down a constant 9 V battery-powered supply. Critically, this lick-detection circuit is optically isolated from the TTL circuit that reports lick events to Teensy, which avoids crosstalk and electrical noise that a shared circuit would introduce.

**Figure 1.**
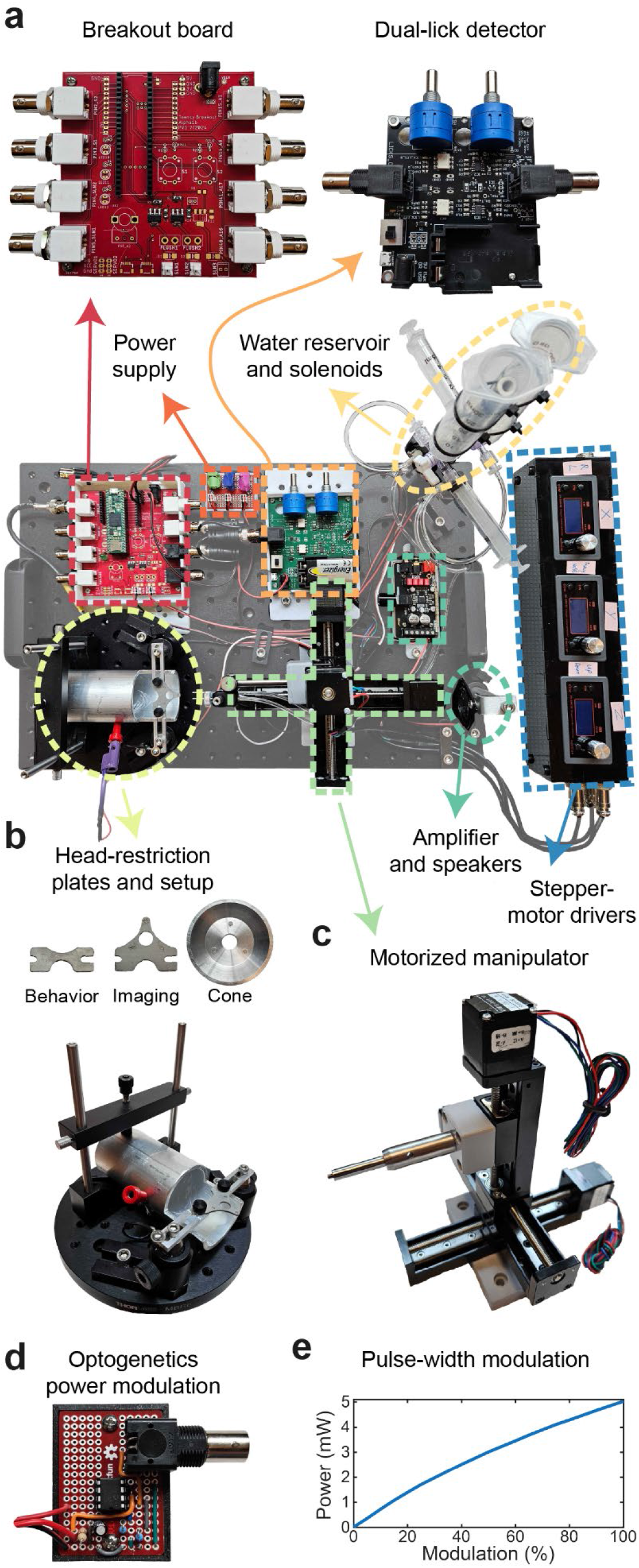
Hardware for head-restrained decision-making tasks in mice. (A) Representative breadboard assembly showing the platform’s principal subsystems: a custom Teensy breakout board with integrated solenoid drivers (“Breakout board”), a conduction-based dual-lick detector (“Dual-lick detector”), a regulated power supply (3.3/5/12 V), a water-delivery subsystem that delivers rewards through solenoids fed by reservoir syringes through 3-way valves (“Water reservoir and solenoids”), a auditory-cue subsystem (“Amplifier and speakers”), and a three stepper-motor drivers used to position the spouts manually or by TTL input (“Stepper-motor drivers”). (B) Two headplate designs: one for behavior (exposed skull for optical-fiber implantation for optogenetics or fiber photometry) and one for imaging (a center hole for a cranial window or GRIN lens). The magnetic water cone mates to the imaging headplate for two-photon water-immersion objectives. Both headplates fit the grounded head-restrain clamp-and-tube assembly to restrict the mouse and complete the lick-detection circuit (bottom). (C) The three-axis motorized spout manipulator composed of three linear tracks driven by stepper motors. (D) The optogenetic power-modulation circuit: an operational-amplifier stage scales Teensy’s native 3.3 V output to 5 V, and Teensy’s pulse-width-modulated (PWM) output monotonically commands LED or laser power. (E) Measured relationship between commanded PWM duty cycle and delivered optical power.

Head restriction can be achieved via two custom headplate designs (Figure 1B): a small plate that leaves the skull exposed for optical-fiber implantation for behavior and optogenetic experiments, and a larger plate with a center hole for a cranial window or GRIN lens for imaging, onto which a magnetically attached water cone mates allowing the use of water-immersion objectives. The latter is particularly useful for high aperture objectives and large field-of-view mesoscopic imaging^8^ where common headplates are bulky and can be unstable, thereby introducing motion artifacts. The mouse is held in a grounded aluminum clamp-and-tube assembly mounted on a magnetic base, which both stabilizes the head and completes the lick-detection circuit. Three linear tracks under stepper-motor control (Figure 1C) position the spouts, either manually using the stepper-motor controller display or under TTL control, such that spout position can be easily adjusted within and between sessions.

To control and calibrate optogenetic stimulation, we built a simple operational-amplifier (Microchip Technology MCP6002) circuit (Figure 1D) that scales Teensy’s native 3.3 V logic output to the 5 V required by common LED drivers and acousto-optic modulators and uses Teensy’s pulse-width modulation (PWM) to command power (Supplementary Figure 4). Measuring delivered optical power as a function of commanded PWM duty cycle showed a linear, monotonic relationship (Figure 1E). We use this signal to convert the requested power in milliwatts into a PWM command for optogenetics stimulation. In contrast, other platforms deliver a TTL trigger for optogenetics onset and offset, changing power on an external device (e.g., LED driver). To comply with these setups, we provide a TTL pulse to trigger optogenetics on an external device.

### Software architecture: on-device state machines and a MATLAB interface

Task logic in SPOUT runs entirely on a Teensy microcontroller to ensure the timing of events is reliable and to offload computational demand from the host computer. The platform is general-purpose by design: two state machines together implement 10 distinct tasks, selected from the MATLAB GUI without recompiling firmware or changing hardware (Figure 2A, top). SPOUT2 drives five common two-spout tasks: uninstructed lick-left/lick-right (LL/LR)^22^, instructed LL/LR^5^, two-armed bandit (2ABT)^17,26^, delayed-response (DR)^7,13^, and Pavlovian conditioning^27^ (Figure 2A, bottom). SPOUT1 drives five often used one-spout tasks: go/no-go^28^, stop-signal^29^, fixed-ratio^30^, progressive-ratio^31^, and Pavlovian conditioning^32^. The two SPOUT state machines differ only in spout number and the complexity of transitions between states (Figure 2B, C). The user controls the state machine via a MATLAB GUI which can load a per-task settings file, update task parameters and report performance in real time, enact manual interventions, activate closed-loop optogenetics, and store data in a standardized manner. Because every task is defined by a settings file rather than by firmware, task parameters are easy to save, version, and share between setups and labs. We released a starting library of validated settings files covering all 10 tasks alongside the code. Task-specific transitions (Figure 2B, C, orange and green arrows) and togglable penalty states (pink boxes) can also be combined independently within each task. Thus, the same two state-machines implement well over a dozen distinct task variants.

**Figure 2.**
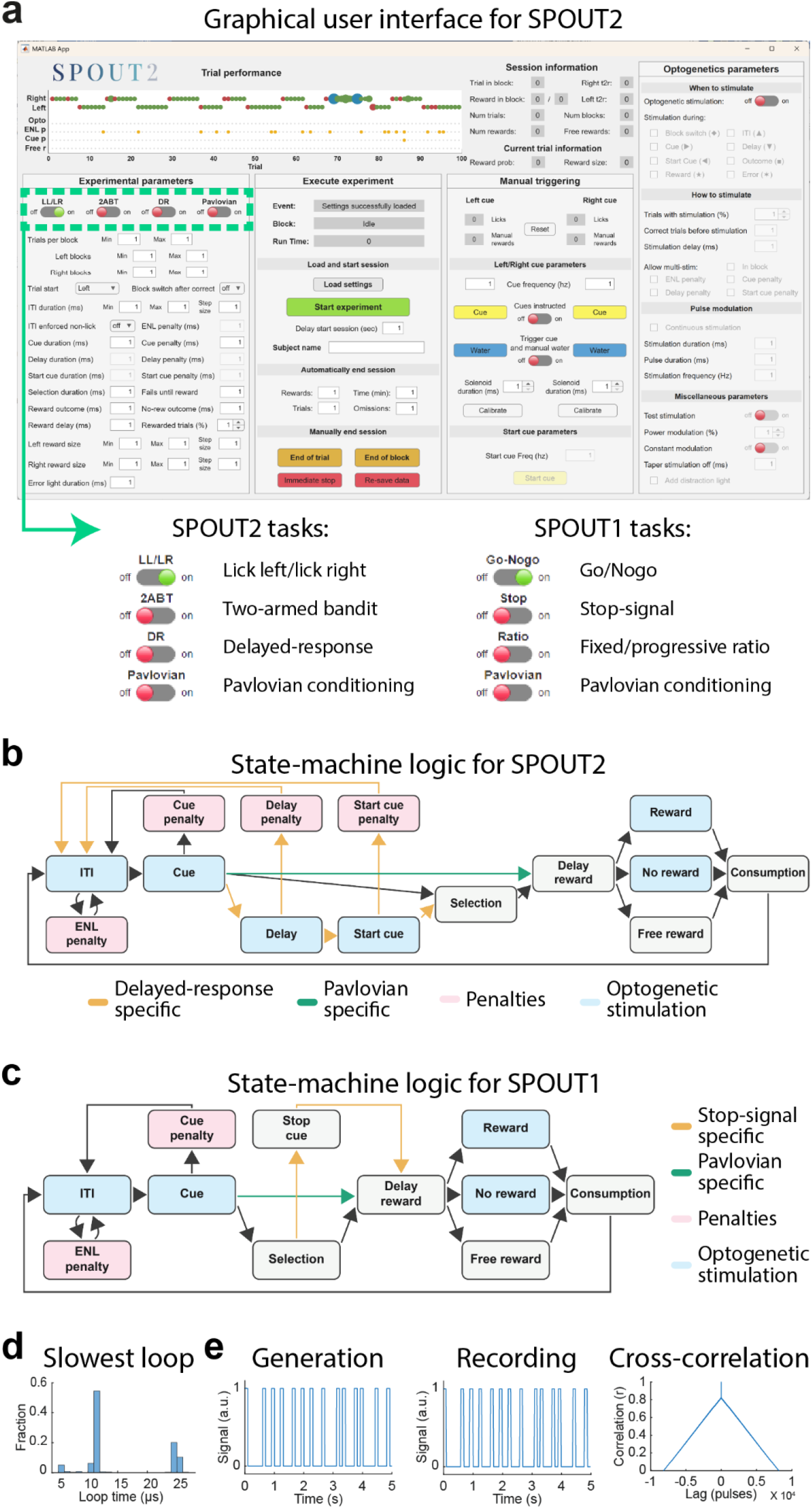
Software architecture: Teensy state machines and the MATLAB graphical user interface. (A) The MATLAB GUI for the two-spout state machine (SPOUT2) loads predefined settings, updates task parameters in real time, reports live task performance, and saves data (top). The same interface toggles task identity (bottom). The two-spout state machine (SPOUT2) implements five common behavioral tasks based on the availability of two spouts: instructed lick-left/lick-right, uninstructed lick-left/lick-right (LL/LR), two-armed bandit (2ABT), delayed-response (DR), and Pavlovian conditioning. The one-spout state machine (SPOUT1) implements five task modes based on the presence of one spout: go/no-go, stop-signal, fixed-ratio, progressive-ratio, and Pavlovian conditioning. The 10 task modes are defined by a shareable settings file and require no additional firmware or hardware, allowing seamless switching between them. (B) State-machine logic for SPOUT2. Gray and blue boxes are states common to every two-spout task (inter-trial interval [ITI], cue, delay, selection, reward/no-reward/free-reward, and consumption); orange transitions are specific to the delayed-response task, green transitions to the Pavlovian task, pink boxes are togglable penalty states (early-lick, cue, delay, and start-cue penalties), and blue boxes mark states in which optogenetic stimulation can be delivered. (C) State-machine logic for SPOUT1, same color code. (D) The slowest sampling period recorded is 26 μs, which corresponds to a sampling rate of 38.5 kHz. (E) SPOUT generates a pseudo-random-interval digital sync pulse (left), which is captured by any external recording system (middle, shown for a benchtop test); cross-correlating the two recorded pulses (pulse duration and corresponding inter-pulse interval) (right) recovers a single, unambiguous alignment at zero lag.

Closed-loop optogenetic stimulation is integrated into the state-machines and the stimulation parameters defining how (e.g., frequency, pulse-width, duration) and when (e.g., restricted epochs, trial number, repeated within blocks) are specified in the GUI. Using the operational-amplifier circuit, precise light power can be set and more complex optogenetics stimulations can be performed such as tapering off stimulation. Epochs are selected through the GUI (Figure 2B, C, blue boxes) and the user can manually validate optogenetics stimulation.

The measured sampling rate of the SPOUT state-machine averaged at 647 kHz ± 0.003 kHz (sampling period of 1.55 μs), while the slowest sampling period recorded was 26 μs, which corresponds to a sampling rate of 38.5 kHz (Figure 2D). This is still ∼5,000 times faster than the 7 Hz lick rate of a mouse^33^, allowing SPOUT to reliably capture licking events. For closed-loop experiments, this means that SPOUT will generally start stimulation 1.55 μs after the trigger event, but certainly within 26 μs, far below any reasonable biological requirement. To align external systems to the SPOUT behavioral timestamps, SPOUT emits a digital synchronization pulse at pseudo-random intervals (Figure 2E, left). This digital pulse (Figure 2E, middle) can be used to recover the correspondence between an external clock and SPOUT’s clock by using cross-correlation on the pulse durations (pulse and inter-pulse interval), which produces a single unambiguous peak at zero lag (Figure 2E, right) rather than an ambiguous or aliased match. In addition, SPOUT can capture digital synchronization pulses from an external system, barring this signal is 3.3 V (Teensy is not 5 V compliant).

### Illustration of a behavioral task: mice learn a history-dependent two-spout LL/LR task

To validate the system, we first examined whether the platform detects individual licks as faithfully as an independent, dedicated recording system (National Instruments analog breakout board controlled by a data acquisition application WaveSurfer). The dual-lick detector is an electrically isolated, conduction-based design in which the mouse is grounded and the spouts pulled up to 9 V. Closing this circuit is sensed and reported as an analog signal to SPOUT and WaveSurfer. Comparing lick times registered by SPOUT against those recorded simultaneously on WaveSurfer from the same analog lick signal, the two systems agreed almost perfectly, with SPOUT registering slightly more licks than WaveSurfer (Figure 3A). This small increase in number of licks was found to be caused by WaveSurfer’s low sampling rate (1 kHz vs 647 kHz): WaveSurfer missed small lick interruptions that SPOUT did accurately record. In a well-trained mouse, licking was tightly locked to the task: mice made a fast choice lick, increased licking after cue presentation, and outcome licks continued on the currently rewarded side within a block (Figure 3B, left). Choice licks and outcome licks each fell in narrow, unimodal windows (Figure 3B, second and third panels), suggesting the system accurately records and does not drop licks. Mice licked at a mean rate of approximately 6 Hz (Figure 3B, right) as previously reported^33^. Together, this indicates that the dual-lick detector resolves licks with the temporal precision needed to analyze behavior at the level of single licks.

**Figure 3.**
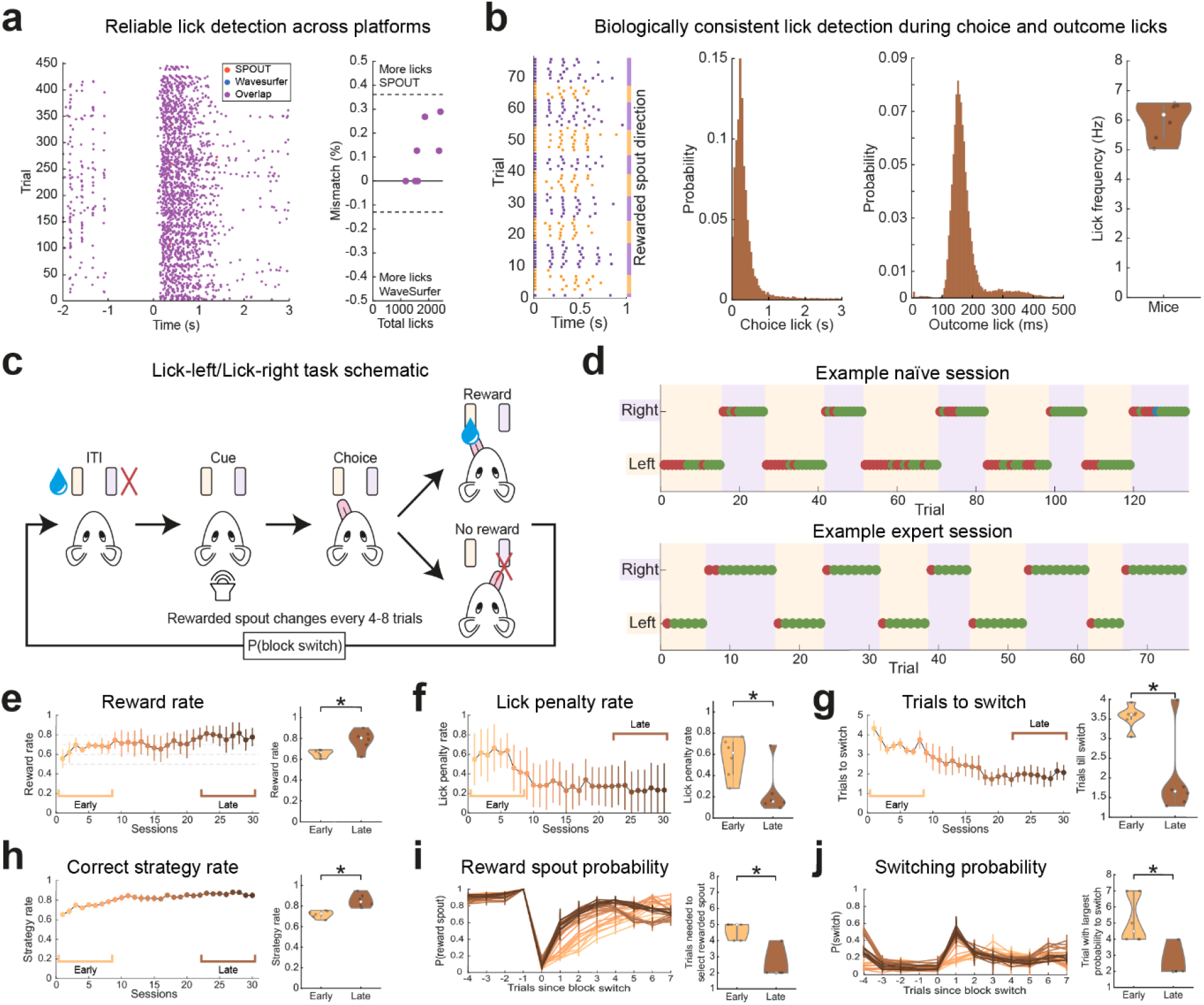
Mice learn a history-dependent lick-left/lick-right (LL/LR) task. (A) Lick signals recorded simultaneously by SPOUT and by an independent data-acquisition system (WaveSurfer) agree almost perfectly (left, raster of lick times recorded by each system across trials); the small excess of SPOUT-detected licks (right, mismatch as a percentage of total licks per session) is consistent with SPOUT’s higher sampling rate. (B) In a well-trained mouse, licking is tightly locked to task structure. Lick raster shows choice and outcome licks tracking the currently rewarded side within a block (left). Choice licks fall in a narrow, unimodal window after cue onset (second panel). Outcome-period licks are likewise unimodal with a rightward skew (third panel). Mice lick at a mean frequency of approximately 6 Hz (right, one point per mouse). (C) Schematic of the LL/LR task: after an ITI in which licking is penalized, an uninstructed auditory cue prompts a choice between a left and a right spout, only one of which is rewarded. The cue itself carries no information about the rewarded side, so the mouse must infer it from its own recent action–outcome history. The rewarded side switches, uncued, after 4–8 non-consecutive rewarded trials (a block). (D) Example naive (top) and expert (bottom) sessions from the same mouse: correct (green) and incorrect (red) choices across trials, colored by block. Naive sessions show gradual drift in choice direction after a block switch; expert sessions show an abrupt switch. (E–J) Behavioral change across 30 training sessions (left of each panel session averages over training; right first eight “early” versus last eight “late” sessions): (E) reward rate increases; (F) the rate of penalized ITI licks decreases; (G) the number of trials needed to select the rewarded spout after a block switch decreases; (H) use of the optimal win-repeat/lose-switch strategy increases; (I) the probability of selecting the rewarded spout, aligned to block transitions, sharpens with training; (J) the probability of switching spouts, aligned to block transitions, sharpens correspondingly. Data plotted as mean ± s.e.m. across animals, n=6 mice; *p < 0.05.

We used SPOUT to train mice on a history-dependent lick-left/lick-right (LL/LR) task, a commonly used task to study decision making in head-restrained mice^22–25^. The task was organized into blocks in which one spout was rewarded, and the other spout was unrewarded. After a variable inter-trial interval (ITI, 1-2 s), during which any lick restarted the ITI, an auditory cue signaled the start of a 3 s selection window, without informing the mouse which spout was rewarded. Selection of the rewarded spout resulted in immediate water reward delivery, while the other spout would not. After pseudo-random 4-8 non-consecutive rewarded trials, the state-machine switched the rewarded spout and the mouse had to update its behavior accordingly (Figure 3C). Naïve mice (early sessions) drifted slowly toward the new side after a block switch, whereas expert mice (late sessions) switched abruptly (Figure 3D).

Across 30 training sessions (n=6 mice), reward rate increased from chance level to approximately 80% (Figure 3E; early vs late t(5)=-3.077, p=0.028), the rate of penalized ITI licks fell (Figure 3F; early vs late t(5)=4.291, p=0.008), and the number of trials needed to select the rewarded spout after a block switch fell toward the one-trial floor (Figure 3G; early vs late t(5)=3.958, p=0.011). Use of the optimal win-repeat/lose-switch strategy, repeating a rewarded choice or switching after an unrewarded one, increased over the same sessions (Figure 3H; early vs late t(5)=-5.186, p=0.004). Aligning choices to block transitions directly showed the same learning: the probability of choosing the rewarded spout dropped sharply on the first trial after a switch and recovered within a few trials, and this recovery sharpened with training (Figure 3I; early vs late t(5)=4.719, p=0.005), as did the corresponding probability of switching sides (Figure 3J; early vs late t(5)=3.576, p=0.016). Together, these results indicate that mice learn to use action–outcome history, rather than the cue, to guide choice in the LL/LR task, and that SPOUT’s behavioral readout is precise enough to quantify that learning trial-by-trial.

### Closed-loop optogenetics: ALM inhibition biases choices ipsilaterally

Anterior lateral motor cortex (ALM) is the canonical lick-planning region in the mouse^7,13,34,35^. Perturbing ALM activity while monitoring behavior provides a validated test of whether SPOUT can deliver precisely timed perturbation and detect a known-direction behavioral consequence in the LL/LR task. We employed VGAT-ChR2-EYFP mice that express channelrhodopsin-2 in inhibitory neurons, which robustly reduces the activity of cortical excitatory neurons upon blue light stimulation^36^. Histology confirmed EYFP expression in ALM (Figure 4A) and fiber-tip placement across 12 hemispheres in 6 mice (Figure 4B).

**Figure 4.**
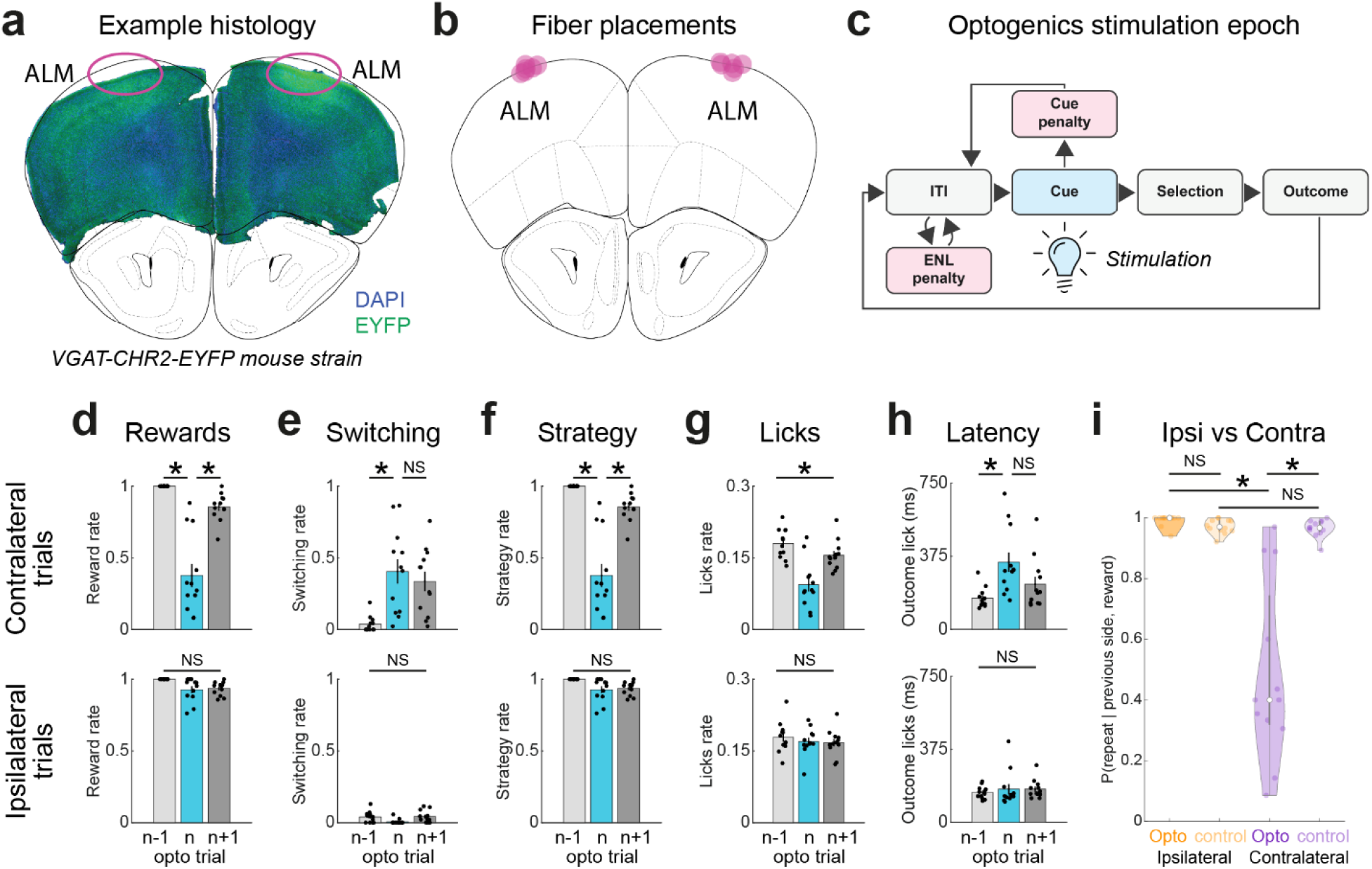
Optogenetic inhibition of anterior lateral motor cortex (ALM) biases mice toward ipsilateral choices. (A) Coronal section from a VGAT-ChR2-EYFP mouse showing bilateral EYFP expression (green) in ALM (outlined) over a DAPI staining (blue). (B) Histologically reconstructed fiber-tip locations in ALM across implanted animals (n=6 mice, 12 hemispheres). (C) Trial-structure schematic showing the closed-loop optogenetic stimulation epoch directly triggered by the state-machine. Stimulation was delivered unilaterally, once per block, beginning at cue onset and terminating at the choice lick or at outcome onset, whichever came first. (D–H) Effect of stimulation on contralateral-block trials (top row) versus ipsilateral-block trials (bottom row, relative to the stimulated hemisphere), comparing the stimulated trial (n) with the immediately preceding (n-1) and following (n+1) unstimulated trials: (D) reward rate drops sharply on the contralateral stimulation trial (n) and recovers on the following trial (n+1), with no effect on ipsilateral trials; (E) switching increases on the contralateral stimulation trial and does not significantly recover on the following trial, with no effect ipsilaterally; (F) use of the correct win-repeat/lose-switch strategy falls on the contralateral stimulation trial and recovers on the following trial (mice abandon a spout that was just rewarded) with no effect ipsilaterally; (G) contralateral licking is reduced on the contralateral stimulation trial and recovers on the following trial, with no change ipsilaterally; (H) latency to the first outcome-period lick (first lick after optogenetics inhibition offset) increases on the contralateral stimulation trial and does not significantly recover on the following trial, with no effect ipsilaterally. (I) Direct comparison of the probability of repeating the previously rewarded side for optogenetic versus control (unstimulated) trials, split by ipsilateral and contralateral block identity. Only contralateral optogenetic trials differ from their control and from the ipsilateral optogenetic condition, showing that ALM inhibition specifically biases mice toward ipsilateral choices. Data plotted as mean ± s.e.m. across sessions, one session per hemisphere, n=12 hemispheres from 6 mice, *p < 0.05, NS=not significant

Stimulation (40 Hz, 10 ms pulse duration) was delivered unilaterally once per block after the second correct trial at cue onset and terminated at choice lick or outcome onset, whichever came first (Figure 4C). This closed-loop, epoch-specific protocol follows directly from restricting optogenetic stimulation parameters in SPOUT, rather than from any external source.

On blocks contralateral to the stimulated hemisphere, optogenetic trials showed a sharp drop in reward rate relative to the immediately preceding (n-1) and following (n+1) unstimulated trials (Figure 4D, top; F(2,10)=65.186, p<0.001; n-1 vs n p<0.001; n vs n+1 p=0.002), increased switching (Figure 4E, top; F(2,10)=18.568, p<0.001; n-1 vs n p=0.025; n vs n+1 p=0.376), reduced use of the correct win-repeat/lose-switch strategy as mice abandoned a spout they had just been rewarded on (Figure 4F, top; F(2,10)=62.186, p<0.001; n-1 vs n p<0.001; n vs n+1 p=0.002), reduced contralateral licking (Figure 4G, top; F(2,10)=9.681, p=0.005; n-1 vs n p=0.06; n vs n+1 p=0.117), and increased latency to the first outcome-period lick (Figure 4H, top; F(2,10)=9.388, p=0.005; n-1 vs n p=0.039; n vs n+1 p=0.116). Every one of these effects was absent during ipsilateral blocks (Figure 4D–H, bottom rows; reward rate F(2,10)=3.753, p=0.061; switching rate F(2,10)=4.037, p=0.052; strategy rate F(2,10)=3.753, p=0.061; ipsilateral licking F(2,10)=0.843, p=0.459; latency outcome lick F(2,10)=1.003, p=0.401). These contralateral deficits were largely transient: reward rate, strategy use, and contralateral licking returned toward baseline on the trial immediately following stimulation (Figure 4D, F, G), whereas the increased switching and lick latency persisted in the following trial (Figure 4E, H). Directly comparing the probability of repeating the previously rewarded choice between optogenetic and control trials, split by block laterality, confirmed that only optogenetics manipulation of contralateral trials differed from control (Figure 4I, F(3,33)=33.157, p<0.001; ipsi opto vs control p=0.326; contra opto vs control p<0.001; ipsi opto vs contra opto p<0.001; ipsi control vs contra control p=0.455). Together, these results confirm that ALM inhibition biases mice toward ipsilateral choices, and highlight SPOUT’s ability to deliver precise and specific optogenetics stimulation.

### Combining with two-photon imaging: ALM population dynamics relate to contralateral choices

Mapping neural activity onto behavior is essential to understand how the brain functions and requires that both recording systems are aligned, a non-trivial process. A pseudo-random synchronization pulse allows for the correction of missing data, identifies glitches in the acquisition systems, and rectifies clock drifts. SPOUT generates a ∼2.2 Hz synchronization pulse with variable inter-pulse intervals that are sent as digital 3.3 V pulses to WaveSurfer. In addition, WaveSurfer stores the frame clock of the computer acquiring calcium imaging frames using ScanImage^37^ on its own clock.

We developed a strategy to synchronize the entire recording chain using the pseudo-random synchronization pulse: SPOUT’s event clock, WaveSurfer’s data-acquisition samples, and ScanImage imaged frames. After identifying and correcting missing samples in the WaveSurfer data stream, using cross-correlation on SPOUT’s pseudo-random synchronization pulses (duration of pulse and corresponding inter-pulse interval) (Figure 5A) with the WaveSurfer-recorded copy of these pulses (Figure 5B) reproduced a zero-lag alignment. Elapsed time on the two systems tracked one another linearly across the full session (∼40 min; Figure 5C). However, a small, consistent clock-drift offset between SPOUT and WaveSurfer (approximately 0.0012%) was present throughout the recording (Figure 5D).

**Figure 5.**
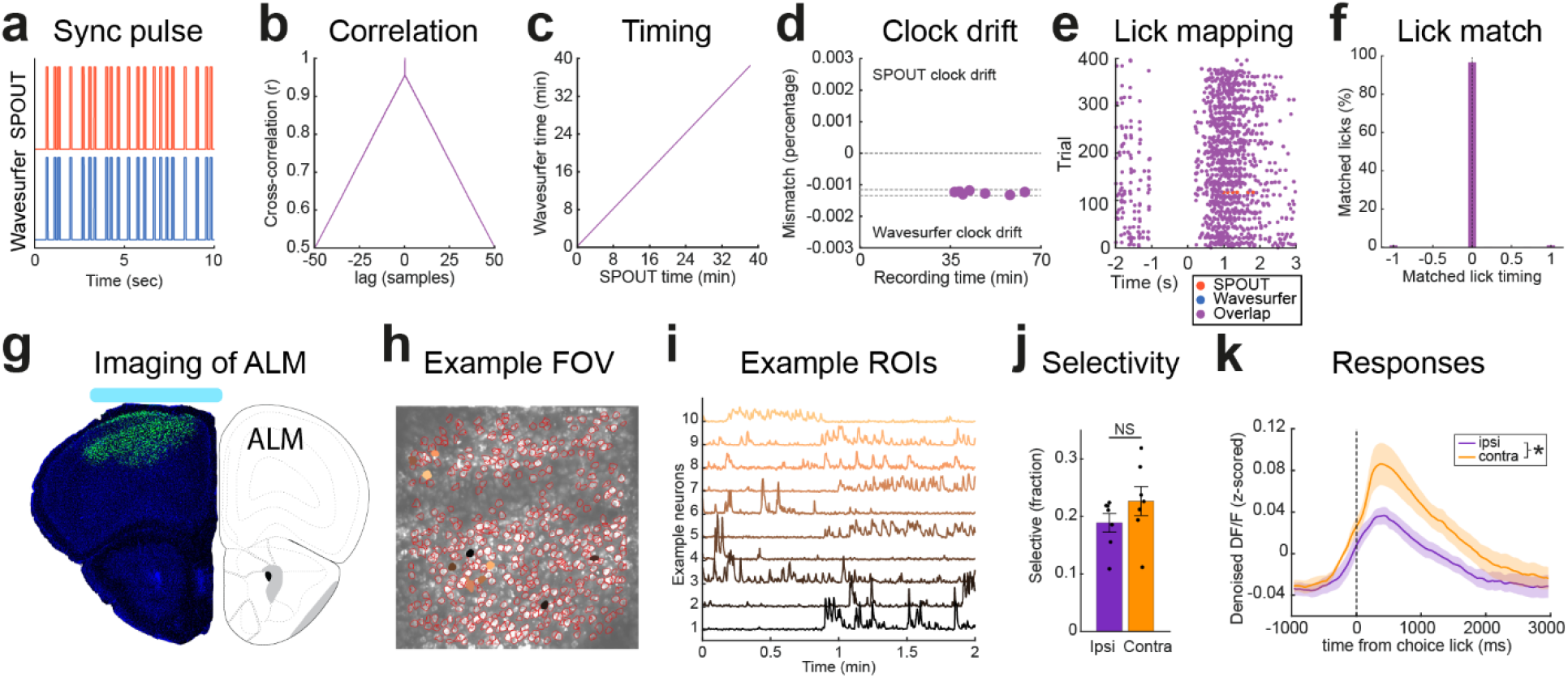
Three-system alignment of behavior, WaveSurfer, and two-photon imaging identifies choice-selective ALM activity. (A) SPOUT’s pseudo-random synchronization pulse (top) independently captured by WaveSurfer (bottom). (B) Cross-correlation of the two recorded pulses (pulse duration and corresponding inter-pulse interval) peaks sharply at zero lag. (C) Elapsed time on SPOUT and on WaveSurfer track each other linearly across a full session (∼40 min). (D) A small, consistent clock-drift offset between the two systems (approximately 0.0012%) is present throughout the recording and is corrected for before downstream alignment. (E) After drift correction, individual SPOUT-detected licks (pink) map onto WaveSurfer time (purple) essentially one-to-one across trials. (F) All licks (100%) are matched within an alignment tolerance of 1 ms, and 96% of all licks match exactly. (G) Histological validation of riboL1-GCaMP8s expression in ALM (left) and its location on the standard atlas (right). (H) Example imaging field of view with segmented ROI contours. Colored ROIs match example traces in I. (I) Example z-scored DF/F fluorescence traces from ten simultaneously imaged neurons. (J) The fraction of choice-selective ALM neurons does not differ significantly between ipsilateral- and contralateral-preferring populations. (K) Population-averaged, choice-aligned responses are larger for contralateral than for ipsilateral choices. Data plotted as mean ± s.e.m. across animals, n=7 mice, *p < 0.05, NS=not significant).

Such offset could accumulate silently over a session and result in severe misalignment later in the session. We corrected for this drift by mapping each synchronization pulse in SPOUT time onto each pulse in WaveSurfer sample space. This generated a lookup table that allowed us to map each SPOUT-detected event directly onto a WaveSurfer sample (Figure 5E). This approach produced a perfect one-to-one match of SPOUT and WaveSurfer licks, with more than 96% of the matched licks coinciding with sub-millisecond precision (Figure 5F). The remaining 4% of the licks matched with less than 1 ms difference. Finally, we used ScanImage frame clock TTL to synchronize calcium imaging frames with behavioral events.

Large field-of-view microscopes with big aperture water-immersive objectives require considerable cones to hold water during imaging sessions^8^. Cones are often glued to the headplate during surgery, resulting in sizable headplates. These headplates often become instable because of their length, resulting in motion artifacts during imaging. Alternatively, they become too thick, resulting in reduced imaging depth. We overcame these limitations by designing a headplate with a small footprint, pressed micro magnets into the headplate, and mounted a magnetic cone exclusively during imaging sessions using a thin layer of silicone to seal the connection (Figure 1B). Using this setup, we imaged and analyzed layer 2/3 excitatory ALM neurons’ activity (n=7 animals) during the LL/LR task, by expressing the calcium indicator jGCaMP8s^38^ in a soma-targeted manner^39^. We confirmed expression and window placement histologically (Figure 5G). An example field of view (Figure 5H) and single-neuron fluorescence traces (Figure 5I) show that individual neurons can be segmented, and their activity tracked across a session. Across imaged neurons, the fraction of neurons classified as choice-selective did not differ significantly between ipsilateral- and contralateral-preferring populations (Figure 5J; ipsi vs contra t(12)=-0.975, p=0.349), but population-averaged, choice-aligned responses were significantly larger for contralateral than for ipsilateral choices (Figure 5K; ipsi vs contra t(12)=-2.219, p=0.047), consistent with previously described findings in ALM^7,13,34,35^. Together, these data suggest that ALM neurons are preferentially activated during contralateral licks and demonstrate that the synchronization pulse of SPOUT guarantees precise timing of events that allow recovery of trial-aligned neural activity.

## DISCUSSION

SPOUT is a versatile, open-source hardware and software platform to study decision-making in head-restrained mice during closed-loop manipulation or recording of neural activity. Its general-purpose framework of two state machines together holds 10 commonly used, unique tasks, which range from classical conditioning to history-dependent two-choice tasks. By changing user-friendly task variables, this can be extended to a dozen or more tasks. SPOUT is user friendly: it does not require programming knowledge and little experience building behavioral equipment. Most of the hardware components are built with off-the-shelf components and require no specific expertise. We provide 3D print files to make some of the components, as well as completed assembly files to outsource constructing technical components. By employing shareable setting files rather than compiling firmware, users can easily share task details, start different experiments without needing to change firmware, and keep track of exact task parameters. In addition, the pseudo-random synchronization pulse allows matching the timing of behavior with external recording systems, despite missing data and clock drifts. Importantly, we show that SPOUT operates on a sub-millisecond regime and detects all licks. Finally, SPOUT’s native closed-loop optogenetics stimulation protocol allows for the design of numerous experiments to examine how brain activity causally relates to behavior.

Microcontroller-based state machines have become the standard in controlling behavioral tasks in modern neuroscience^2,4,17,18,40–43^. With the recent developments in microcontrollers, many Arduino-based systems are being upgraded to Raspberry Pi-based or Teensy-based systems instead to accommodate faster event detection, closed-loop control, and local parameter storage. Previously published systems often solve a single limitation: affordable hardware, flexible and user-friendly software, a variety of behavioral tasks, optogenetics control, or synchronization with other systems. To overcome this limitation, we introduce SPOUT, an open-source, Teensy-based generalized platform that integrates many commonly used, validated decision-making behavioral tasks, off-the-shelf and low-cost hardware, fast sub-millisecond dual-lick detection, integrated and calibrated closed-loop optogenetics control, and a validated framework to align complex behavior with external neural recording devices. By offloading the firmware to Teensy, users can employ relatively cheap and simple computers to run the GUI, without compromising timing, sampling rate, or task efficiency. We provide a starting library of validated settings files for 10 commonly used behavioral tasks, which helps standardize examining behavior in rodents and allows sharing tasks across laboratories. In addition, many more task variants can be configured by changing task variables through our user-friendly MATLAB GUI, without any programming experience required. The hardware to run these tasks is developed and built from off-the-shelf components (e.g., motorized linear tracks to control spout position) or custom-made where the full fabrication design files (e.g., PCB Gerber zip files for dual-lick detectors) are openly shared and provided. These files can be sent to any general PCB fabricator to place an order for assembly by the user (i.e., basic soldering).

Alternatively, we provide full assembly files that allow commercial assembly without any user assemblage. Tasks are organized around lick detection, and our conduction-based, electrically-isolated lick detection system records licks with sub-millisecond latencies, allowing for detailed lick-structure examination. To probe causal relationships between brain activity and behavior, SPOUT allows for closed-loop, detailed optogenetics stimulation based on task state (e.g., cue presentation), animals’ behavior (e.g., licking), or action consequences (e.g., reward outcome). Optogenetics can be delivered at various epochs, continuously or pulsed, at variable intensities (directly controlled by SPOUT or an external device), and tapered off. This level of control will help design better experiments that are tailored to physiological neural activity. Finally, we provide a framework for synchronizing SPOUT behaviors with external recording devices. Previous methods require the behavioral setup to send TTL’s for each behavioral event to a data-acquisition system, limiting the number of task-related signals that can be captured. In addition, any hiccup or glitch in the data-acquisition system would result in immediate loss of data and potentially loss of the rest of the session when neural data is acquired at a fixed sampling rate. We show and provide scripts to align the raw behavior output to a data-acquisition system despite system glitches and clock drift. This will allow users to combine SPOUT with their preferred neural recording system and behavioral cameras, while maintaining synchronized streams of data and rich datasets. Taken together, SPOUT provides an affordable platform of hardware, software, and analyses code that integrates all requirements expected from a modern neuroscience experiment.

Accurate lick detection in mice is challenging and requires dedicated hardware. Most common means of detecting licks revolve around optical-, force-, and conduction-based designs^44–46^. Optical lick sensors usually require the tongue to interrupt an infrared beam shortly before the drinking spout is reached. This type of sensor often fails to distinguish licks in a bout, resulting in an under-estimation of the behavior of the mouse^47^. In addition, dissociating left versus right licks is challenging when spouts are close together. Force-based lick sensors require an animal to physically press a spout with enough force to trigger a lick recording. Generally, these sensors have different sensitivities depending on the volume of liquid presented to the animal^47^. Conduction-based sensors are commonly used and require the animal to close a small electrical circuit^45^. A common problem with commercially available conduction-based sensors is the production of significant electrical artifacts, resulting in spurious licks as well as distorted neural signals. To overcome this limitation, we optically-isolated the detection circuit from the output stage, preventing crosstalk between detection and reporting of licks. To reliably detect licks, the system needs to run at a high sampling rate to 1) detect each lick event and 2) precisely identify the onset and offset of a lick. Previous systems sample with supra-millisecond sampling rates^17,40,48^, potentially missing licks. We make use of the rapid clock speed of Teensy and optimized our state-machines for fast sampling rates.

Therefore, we report an average sampling rate of 647 kHz, which corresponds to a sampling period of 1.55 μs. The slowest sampling period recorded with SPOUT was 26 μs, which exceeds other methods in terms of speed^2,4,17,40,48^. Together, our optically-isolated, conduction-based lick detector allows reliable detection of licks and resolves lick onsets and offsets.

To highlight the capabilities of the system, we trained mice on a history-dependent LL/LR task where mice have to use action-outcome associations to successfully earn rewards^22–25^. Like previous studies, we found that mice use recent reward history to update their behavior^22^. We used the closed-loop optogenetics capabilities of SPOUT to deliver precisely timed perturbation to the ALM, a region critically involved in planning of lick behavior in the mouse^7,13,34,35^. We found that briefly inhibiting ALM during cue presentation strongly disrupts contralateral performance by inducing switching behavior. These effects were absent during ipsilateral trials, emphasizing the lateral nature of ALM and the specificity of SPOUT. Finally, SPOUT generates a pseudo-random synchronization pulse, which can be used to align behavior to neural recordings post-hoc. We show that this pulse can identify recording glitches and overcome clock drifts, which can result in critical alignment complications. Using a novel, stable headplate for large field-of-view microscopes, we recorded calcium dynamics in ALM using two-photon microscopy. Although ALM contains an equal distribution of ipsilateral and contralateral preferring neurons, population activity is more strongly recruited during contralateral trials. These results suggest that ALM neurons drive contralateral licking through enhanced population dynamics, which can be abolished by optogenetics inhibition. Together, these results indicate that SPOUT can be used to train complex decision-making tasks in mice, provides closed-loop and temporal precise optogenetics perturbation, and can be employed together with high-dimensional neural recordings.

The current SPOUT platform has several limitations, which can be overcome for certain experiments. First, the platform is built for head-restrained mice. However, with few hardware adjustments, it is possible to study freely behaving mice or other laboratory animals. In addition, choice behavior is required to be made by licking a lick spout. For certain experiments, the user might want to dissociate the reward delivery from the choice modality (e.g., pushing a lever). The firmware simply requires a changing signal, which means that any electrical switch could function as making a “choice”. Next, the user interface requires MATLAB software. We do not see this to be a limitation for adopting SPOUT as MATLAB is still widely used in neuroscience research and licenses are often provided by universities and research facilities. Finally, SPOUT is currently limited to 10 commonly used behavioral tasks. By changing global task settings in the GUI of SPOUT, users can define many more task variants without any programming experience. Users with programming experience can integrate new behavioral tasks. In addition, SPOUT is available on Github, and future releases might provide more behavioral tasks.

The SPOUT platform is a versatile, open-source hardware and software platform that is well-validated and easy to implement by new users and new labs. The system is scalable, flexible, and compatible with many external devices. By employing off-the-shelf components and full assembly files, we balance affordability and effortless implementation. The platform is modular and components can be easily adjusted, swapped, or upgraded. In addition, the system is highly consistent and durable as we have not replaced a single component across thousands of behavioral sessions. Taken together, SPOUT is a user-friendly and flexible platform for diverse neuroscience experiments, can be easily adopted with little hardware or programming knowledge, provides temporally precise closed-loop manipulations, and allows easy integration with in vivo neural dynamics recordings.

## MATERIALS AND METHODS

### Animals

All procedures were approved by the Harvard Standing Committee on Animal Care and conducted in accordance with the US NIH Guide for the Care and Use of Laboratory Animals. Experiments used adult male mice, group-housed under a reversed 12:12 h light:dark cycle and tested during the dark (active) phase. Male mice aged 8 weeks to 4 months were used. Behavioral training used 6 mice VGAT-ChR2-EYFP^36^ mice (heterozygous; bred in-house, originally JAX 014548). The same 6 mice underwent optogenetic inhibition of anterior lateral motor cortex (ALM). Two-photon imaging of ALM was performed in a cohort of 7 vGlut1-Cre mice (heterozygous; bred in-house, originally JAX 023527). All behavioral animals were group-housed and water restricted to 85% of their body weight, with free access to food and no access to water. Animals earned water during daily training but received supplemental water if daily task-earned volume fell short.

### Hardware

The behavioral monitoring and control system (Figure 1A) is built around a custom designed Teensy breakout board with integrated solenoid drivers (5 V, Supplementary figure 2), a conduction-based dual lick detector (3.3 V, Supplementary figure 3), an audio amplifier (12 V, Drok 15 W * 15 W) and speakers (Intervox ICC 450 S150RMSS 0.2 W) for auditory cues, and three linear tracks driven by stepper motors (Befenybay BE073-1 with NEMA11) and controlled by three stepper-motor drivers (Banria 4257). All equipment is mounted onto the breadboard using custom-designed 3D printed housing. The motor drivers allow for positioning the spouts in front of the mouse manually or by TTL input (Figure 1C). Reward water is gated by silent solenoid valves (5 V, The Lee Company LHQA0531220H) fed from reservoir syringes (30 ml) and cleared through flush syringes (5 ml) via 3-way valves. The volume delivered per reward is set by the calibrated open duration of each spout’s independent solenoid, which we daily calibrated to dispense 2.5 uL water per reward.

Licks are detected by a battery-powered, electrically isolated, conduction-based circuit. Contact between the grounded mouse and a stainless-steel spout induces a voltage drop on the spout, which is pulled up to 9 V through a 4.7 MΩ resistor. The spout voltage is sensed and compared against a user-adjustable threshold. The input stage utilizes a low-pass RC filter with a time constant of 10 microseconds and employs 100 mV of hysteresis. The thresholded signal is optically coupled to a 3 V output stage consisting of a line driver and a 50 Ohm series source terminator to drive coaxial loads. The digital lick event is reported to Teensy via a 50 Ohm coaxial BNC cable. For imaging experiments, this digital lick signal is split and a copy sent to an analog acquisition board (National Instruments PCIe-6341) with a BNC breakout board (National Instruments BNC-2090A) recorded by a data acquisition application (WaveSurfer 1.0.6), which allows for directly comparing licks detected by SPOUT and an independent data-acquisition system. The animal is restrained in a grounded aluminum clamp-and-tube head-restraint assembly (Figure 1B) mounted on a magnetic base, which allows for convenient mounting and unmounting of the animal and completes the lick-detection circuit. During imaging sessions, a magnetically attached water cone mates to the imaging headplate and was sealed with a thin layer of silicone (Picodent Eco-sil speed) to hold water for the two-photon objective.

Optogenetic stimulation power is set by a custom operational-amplifier circuit (Figure 1D) that scales Teensy’s native 3.3 V logic output to the 5 V required by common LED drivers and acousto-optic modulators and uses Teensy’s pulse-width modulation (PWM) output to command power linearly. PWM changes the duty cycle resolution in 8-bit (0-255). We calibrated this circuit before each stimulation session by measuring delivered optical power at a series of commanded PWM duty cycles with a photodiode power meter (Thorlabs PM100D). This allowed us to convert the requested power in milliwatts to a PWM percentage command during optogenetics stimulation.

### Firmware and software

The firmware runs on a Teensy 4.1 microcontroller (600 MHz ARM Cortex-M7). Sampling timing was monitored online by recording the number of script loops per second bin and the maximum sampling period within that bin over the course of sessions. The main sampling rate averaged at 647 kHz (± 0.003 kHz), which corresponds to a sampling period of 1.55 μs. More importantly, the slowest sampling rate recorded was 38.5 kHz, which corresponds to a sampling period of 26 μs. Running the firmware on Teensy allowed us to reach such high sampling rates and freed up performance on the host computer. We developed two state machines: SPOUT1 for one-spout tasks and SPOUT2 for two-spout tasks (Figure 2B, C). The difference between SPOUT1 and SPOUT2 is the different state transitions. Each state machine is a fixed transition of states (inter-trial interval, cue, delay, selection, reward/no-reward/free-reward, consumption, and togglable penalty states) whose active transitions are set by the task mode selected in the companion MATLAB GUI (App Designer, Figure 2A). Switching task identity requires no recompiling and no hardware change, only a different settings file. SPOUT1 supports five common task modes: go/no-go^28^, stop-signal^29^, fixed-ratio^30^, progressive-ratio^31^, and Pavlovian conditioning^32^. SPOUT2 supports five common tasks: instructed^5^ and uninstructed lick-left/lick-right^22^, two-armed bandit^17,26^, delayed-response^7,13^, and Pavlovian conditioning^27^. Together SPOUT1 and SPOUT2 provide 10 commonly used decision-making tasks. We employ the EEPROM of Teensy 4.1 to store last task settings, even when Teensy is powered off.

The MATLAB GUI requires an installation of MATLAB (tested on version 2021b till 2026a). The GUI can load the task settings from a CSV file that is generated per-task and per-animal, which can be edited, versioned, and shared between setups and labs like any other text file. After each session, the GUI stores an updated task setting file for future references. We release a starting library of validated settings files for all 10 modes alongside the rest of the code, so a new user can reproduce a given task based on common sets of parameters. Every session writes a set of human-readable CSV files which contain settings, behavioral data (all events, block averages), and synchronization information (internally generated and externally recorded). The GUI allows task parameters to be updated in real time during a session, plots live task performance, and configures which state-machine epochs are eligible for optogenetic stimulation. In addition, the user can change subject name, delay the start of the session, manually or automatically end a session (based on number of rewards, trials, omissions, or time), and manually provide cues and rewards. Optogenetics stimulation is temporally precise and can be fully customized by the user, allowing for precise control over when and how to stimulate. Stimulation epochs include at a block switch, during ITI, cue, delay, start cue, outcome, reward, or error. We included ways to manually test stimulation, taper/ramp stimulation off, and modulate the power of stimulation.

Finally, the state-machines generate a constant synchronization pulse (∼2.2 Hz, 100 ms pulse duration, random inter-pulse interval of 110-590 ms) starting when the task initiates and stored in a CSV file (*_SyncPulses.csv). This synchronization is a digital 3.3 V pulse that can be recorded by peripheral systems, such as WaveSurfer, to allow post-hoc alignment of data. In addition, if the user prefers to record an external synchronization pulse (max 3.3 V as Teensy is not 5 V compliant), the state-machine will store this (*_ExtSyncPulses.csv) as well. It is important to generate a random synchronization pulse to identify data acquisition system glitches and dropped packets, which cannot be determined by a static signal.

### Surgery and headplate implantation

Mice received carprofen in their drinking water (66 ug/ml) at least one day before until three days after surgery. Animals were anesthetized with isoflurane (4% induction, 1-1.5% maintenance) and regular air under aseptic conditions using a stereotaxic frame (David Kopf Instruments Model 1900). During surgery, eyes were covered with eye lube (Optixcare) and temperature maintained at 37 °C. All reagents were injected with a custom-build system using a syringe (Hamilton 5 μl) mounted in a pump (World Precision Instruments, 78-8110W) and tubing connecting a pulled (Sutter Instrument Co. P-97) glass pipette (Drummond Scientific Company pipettes) with a tip of approximately 50 μm. To improve targeting, the pipette was slowly lowered into the brain (500 nm/min) 100 μm ventral to the target region before being brought up to the injection location. Reagents were injected (100 nl/min) and the pipette was left in place for 5 min. Pipette was slowly retracted to 100 μm dorsal to the target region before being removed from the brain. After surgery, mice recovered in a heated home cage.

For optogenetic experiments, VGAT-ChR2-EYFP mice, which express channelrhodopsin-2(H134R)-EYFP under the vesicular GABA transporter promoter and require no viral injection, received bilateral 400 μm craniotomies above the ALM (AP +2.5mm, ML +/- 1.5 mm) made with an electric drill and a ball bur (FG 1/4 Wave Dental). Mice received bilateral chronic optical fibers (200 μm core, 0.37 NA, 1.25 mm ferrule; Neurophotometrics) targeting ALM and implanted above the dura. Fiber locations were marked with crimson fluorescent microspheres (FluoSpheres, 10 μm; ThermoFisher) injections. Fibers were secured to the skull with cyanoacrylate glue (Loctite 454), and a custom titanium behavioral headplate (Figure 1B) was amounted on the cerebellar plate and secured with dental cement (C&B Metabond).

For imaging experiments, vGlut1-Cre mice received, in addition to carprofen water, dexamethasone (6.5 mg/kg, to reduce brain edema), buprenorphine (0.6 mg/kg, opioid analgesia), Baytril (60 mg/kg, antibiotic), and Ketofen (6 mg/kg, anti-inflammatory). A 2 mm craniotomy was drilled over the left-hemisphere ALM (centered at AP +2.5 mm, ML 1.5 mm) and this part of skull removed. Dura was removed and a Cre-dependent, soma-targeted calcium indicator, AAV5-DIO-RiboL1-GCaMP8s (titer 3.62 x 10^12 vg/ml), was injected into ALM as eight 200 nl boluses spaced across the 2 mm region at DV -0.7 mm. A custom chronic imaging window fabricated from laser-cut coverslips (Avator 48393-106 No.1; two stacked 2 mm and a 3 mm coverslip glued together with Norland 81 optical glue) was then seated over the craniotomy and gently pushed down. The window was glued to the edged skull using cyanoacrylate glue (Loctite 454), the skull covered with dental cement (C&B Metabond), and a custom titanium imaging headplate (Figure 1B) secured to the skull.

### Behavioral task and training

Mice were trained on a SPOUT2 task: history-dependent lick-left/lick-right (LL/LR) task which is a commonly used head-fixed dual-spout procedure^22^. Each trial began with an enforced no-lick inter-trial interval (ITI) drawn uniformly from 1.0, 1.25, 1.5, 1.75, or 2.0 s distribution, during which any lick restarted the interval. If the mouse licked during the single uninstructed 75 ms auditory cue, uninformative in directing which spout is rewarded, the trial was reset. The auditory cue signaled the start of a 3 s selection window, during which the mouse could lick either the left or right spout or not at all, which would result in an omission trial. Importantly, only one spout was rewarded, and a correct choice delivered a water reward with 100% reward probability. A choice lick was followed by a fixed 3 s outcome epoch that occurred whether or not reward was delivered. The rewarded spout switched, without any cue signaling the change, after 4-8 non-consecutive rewarded trials on the same side (i.e., a block).

Before task training began, mice were habituated to handling and head fixation: two days of hand-held water delivery by syringe, three days of water delivery by syringe while head-fixed in the setup, and two sessions of manually alternated left/right rewards delivered through the spouts. Block length was then shaped over training, beginning with blocks of 8, then 6-7, then 4-5, and finally 4-8 rewarded trials (the expert and experimental structure); throughout training, a free reward was delivered after six consecutive unrewarded trials to encourage switching to the rewarded spout. Mice were trained daily on the LL/LR task for approximately a month, with one session per day. Each session ended after 1 h or 40 consecutive omissions, whichever came first. Optogenetics manipulations and calcium imaging began only after mice reached expert performance (>2 consecutive sessions of 80% reward).

### Optogenetics

In VGAT-ChR2-EYFP mice implanted with bilateral ALM fibers, optogenetic stimulation was delivered unilaterally (only one hemisphere per session) on the third rewarded trial of each block (approximately 15% of trials), and was restricted to a single state-machine epoch that began at cue onset and ended at the first choice lick or after 3 s (the duration of the selection window), whichever occurred first. Light was provided by a 473 nm DPSS laser (Laser Quantum Ventus + MCP 600), its power modulated by an acousto-optic modulator (AA Opto-Electronic MTS110-A3-VIS) driven through the PWM-calibrated command circuit described above and gated by a mechanical shutter (Vincent Associates), and delivered as 40 Hz trains of 10 ms pulses at on average 6.9 mW (range 3-10 mW) at the fiber tip across the stimulation epoch. Output power was calibrated at the fiber tip before each session with a power meter (Thorlabs PM100D). Trials were classified as contralateral or ipsilateral relative to the stimulated hemisphere and analyzed against the immediately preceding (n-1) and following (n+1) unstimulated trials of the same block type.

### Two-photon calcium imaging

Imaging was performed through the chronic window over left-hemisphere ALM on a custom two-photon random-access mesoscope^8^ controlled by a computer running ScanImage (2023)^37^, with excitation at 920 nm from a tunable Ti:sapphire laser (Coherent Chameleon) and approximately 78 mW excitation light at the sample. ScanImage sends a digital acquisition TTL to WaveSurfer (i.e., high signal for the duration of acquisition). Layer 2/3 of ALM was imaged over a 500 x 500 μm field of view (1000 x 1000 pixels) at 21.47 frames per second. Before each session the window was cleaned, a thin layer of two-component silicone (Picodent Eco-sil speed 2) was applied to the headplate, a water well was lowered onto it and filled with deionized water, and the objective was lowered into the well. Raw movies were examined in FIJI^49^, motion-corrected and segmented into regions of interest (ROIs) using Suite2p^50^, manually curated in PRISM, and denoised and deconvolved with OASIS^51^.

### Histology

After the final session, mice were transcardially perfused with phosphate-buffered saline (PBS) followed by 4% paraformaldehyde (PFA). The head was post-fixed in 4% PFA overnight, after which the brain was removed, cryoprotected in 30% sucrose for two days, and sectioned at 50 μm on a cryostat (Leica CM3050S). Sections were immunostained for the reporter with rabbit anti-GFP (ThermoFisher Scientific A11122) and an Alexa Fluor 488 goat anti-rabbit secondary (ThermoFisher Scientific A11008), and counterstained with DAPI (Sigma-Aldrich D9542). Sections were mounted, imaged on a slide scanner (Olympus VS200), and processed in QuPath to localize fiber placements and injection sites in ALM.

### Multi-clock synchronization and data alignment

Behavioral data from SPOUT, an independent data-acquisition system (WaveSurfer 1.0.6, Janelia Research Campus), and two-photon imaging frames were reconciled onto a common timeline as follows. SPOUT generates and timestamps a digital synchronization pulse with a fixed 100 ms high phase and a pseudo-random 110–590 ms low phase. This pulse is recorded as a digital channel by WaveSurfer during every session. Matching pulse-to-pulse intervals between the two recorded copies of this signal detects any WaveSurfer samples dropped during acquisition (gaps), which are patched (NaN-filled) before further processing. Every successfully matched sync pulse becomes an anchor point associating a SPOUT timestamp with a WaveSurfer sample index. Anchors are stored per session and absorb clock drift between the two systems rather than assuming a single fixed offset, since we found a small, consistent WaveSurfer clock drift of approximately 0.0012% across a session. Any SPOUT event timestamp (e.g., a lick, cue onset, or choice) is then placed in WaveSurfer-sample space by linear interpolation between its two nearest anchors. Two-photon imaging frames are mapped onto WaveSurfer time independently, using each frame’s known acquisition duration to bridge the same WaveSurfer gaps. For each trial, imaging frames between that trial’s start and stop times (in WaveSurfer-sample space) are extracted, and both the behavioral event fields and the per-frame acquisition times are re-expressed in milliseconds relative to that trial’s own cue onset, producing one cue-aligned trial per entry in the trial-level data structure used for all imaging analyses. To align imaging data to an arbitrary behavioral event (e.g., a choice or outcome lick) for plotting, we built a common output time axis from the per-session imaging frame rate and the desired pre/post window, shifted each trial’s frame times by that trial’s own event time, and, for every point on the common axis, assigned the nearest real frame within half a frame period. A bin was left as a missing value if no real frame fell within that tolerance. This nearest-frame matching is used to build every trial-by-trial imaging array in this study. Calcium traces themselves are never resampled or interpolated, only the correspondence between a desired time point and the nearest acquired frame. Alignment quality was quantified by cross-correlating SPOUT and WaveSurfer copies of the sync signal pulses, by comparing elapsed SPOUT and WaveSurfer time across the session, and by mapping individual SPOUT-detected licks onto WaveSurfer time and computing the fraction that coincided with an independently detected WaveSurfer lick within the alignment tolerance. Importantly, this synchronization method is robust against acquisition system hiccups, glitches, and clock drifts. In addition, it is not restricted to two-photon calcium imaging and can be used for various techniques such as electrophysiology or one-photon miniscope imaging^52^.

### Choice selectivity and event-aligned activity

All calcium imaging analyses used the z-scored denoised calcium trace. For each trial, activity was aligned to the choice lick and extracted over a window of 1 s before to 3 s after this event. To identify choice-selective ROIs, we compared each neuron’s mean response on ipsilateral (left) versus contralateral (right) correct-choice trials over the 500 ms post-choice window using an independent-samples t-test. A neuron was classified as selective if p < 0.05, and was labelled ipsi-or contra-selective according to which choice evoked the larger mean response; neurons with fewer than two trials per side were left untested, and sessions required at least five ipsi and five contra correct trials to be included.

Across sessions, population activity is plotted as the mean ± SEM (across sessions) of the neuron-averaged trace for each condition (ipsi-and contra-lateral), interpolated onto a common 30 Hz time axis, with conditions compared by an independent-samples t-test on the mean activity over the first 500 ms after the event.

### Statistical analysis

Statistically comparing early versus late performance used paired t-tests. Analyzing optogenetics-effects used one-way ANOVA, followed by post-hoc tests corrected for multiple comparisons. Number of choice-selectivity between ipsilateral- and contralateral-preferring ALM neurons and comparing their responses used independent t-tests. Statistical significance was set at p < 0.05 (denoted * in all figures; NS indicates p ≥ 0.05). All statistics were computed in MATLAB (R2024b and R2026a).

## Data availability

All firmware, GUI source code, task-settings files — including a starting library of validated settings files covering all 10 SPOUT1/SPOUT2 task modes — preprocessing and analysis code, hardware design files (CAD and PCB), a bill of materials, and assembly and quality-control documentation are available at our GitHub repository (https://github.com/spout-task) under MIT license, with a versioned release archived on Zenodo.

## Acknowledgements

We would like to thank Rebecca N. Alvarado and CT King for help with animal husbandry, Elizabeth Drewry for administrative support, Michelle Ocana for imaging support, and John LeBlanc for technical support. This work was supported by the EMBO fellowship (1288-2024, B.J.G.V.D.B), Dutch Research Counsil Rubicon (019.242EN.013, B.J.G.V.D.B), Helen Hay Whitney Foundation postdoctoral fellowship (B.J.G.V.D.B), NIH BRAIN Initiative postdoctoral fellowship (F32MH125596, A.E.G.), BBRF Young Investigator Award (A.E.G.), Harvard Medical School Mahoney postdoctoral fellowship (A.E.G.), the Howard Hughes Medical Institute (B.L.S.), the Foundation for OCD Research (B.L.S.), and the NIH (R35NS137336, B.L.S.).

