## Supplementary figures for "SPOUT: An open-source hardware and software platform to study decision making while manipulating and recording from neural activity"

| Description | Company | Model | Unit $ | # | Total $ |
|---|---|---|---|---|---|
| **Breakout board** | | | | | |
| Complete assembly | | | 100 | 1.00 | 100.00 |
| 3D printed housing | | | | 1.00 | 0.00 |
| **Power supply** | | | | | |
| Power adapter 3 V | NC | BY0301&BK | 5.57 | 1.00 | 5.57 |
| Power adapter 5 V | MTYTOT | ALT-0503 | 6.99 | 1.00 | 6.99 |
| Power adapter 12 V | SmoTecQ | STQ-12V-2A | 5.95 | 1.00 | 5.95 |
| 3D printed housing | | | | 1.00 | 0.00 |
| **Dual-lick detector** | | | | | |
| Complete assembly | | | 100 | 1.00 | 100.00 |
| 3D printed housing | | | | 1.00 | 0.00 |
| **Water reservoir, solenoids, and spouts** | | | | | |
| Syringe 5 ml | BD Medical | 309646 | 0.18 | 2.00 | 0.36 |
| Syringe 30 ml | BD Medical | 302832 | 0.32 | 2.00 | 0.64 |
| 3-way valve | Cole Parmer | 48523-38 | 1.005 | 2.00 | 2.01 |
| Solenoid | The Lee Company | LHQA0531220H | 65.6 | 2.00 | 131.20 |
| Tubing (ID 1/16, OD 1/8, wall 1/32) | Tygon | E-3603 | 0.28 | 10.00 | 2.80 |
| Stainless steel tube | McMaster-Carr | 5560K72 | 3.9 | 1.00 | 3.90 |
| Pedestal post holder | Thorlabs | PH1 | 8.74 | 1.00 | 8.74 |
| Optical post | Thorlabs | TR6 | 8.66 | 1.00 | 8.66 |
| Cage rods | Thorlabs | ER3-P4 | 7.255 | 2.00 | 14.51 |
| Swivel post | Thorlabs | MSWC | 40.8 | 1.00 | 40.80 |
| Laser cut holder for solenoids | | | | 1.00 | 0.00 |
| 3D-printed holder for spouts | | | | 1.00 | 0.00 |
| **Head restriction set-up** | | | | | |
| Kinematic magnetic base | Newport corporation | BK-3A | 159.64 | 1.00 | 159.64 |
| Pedestal post holder | Thorlabs | PH1 | 8.74 | 1.00 | 8.74 |
| Optical post | Thorlabs | TR6 | 8.66 | 1.00 | 8.66 |
| Adjustable-Height Optics Clamp | Thorlabs | VG100 | 93.34 | 1.00 | 93.34 |
| Aluminum tube | McMaster-Carr | 6061 | 6.75 | 1.00 | 6.75 |
| Banana socket | na | na | 1.28 | 1.00 | 1.28 |
| Head-restriction plate behavior | SendCutSend | Titanium | 1.4 | 1.00 | 1.40 |
| Head-restriction plate imaging | SendCutSend | Titanium | 1.61 | 1.00 | 1.61 |
| Imaging cone | eMachineShop | Aluminum 6061 | 4.5 | 1.00 | 4.50 |
| Magnets plate | K&J Magnetics | 1/16" x 1/32" | 0.14 | 3.00 | 0.42 |
| Magnets cone | The Magnet Baron | 2 mm x 0.5 mm | 0.15 | 3.00 | 0.45 |
| CNC head-restriction holders | | | | 2.00 | 0.00 |
| Breadboard lasercut | | | | 1.00 | 0.00 |
| **Motorised manipulator** | | | | | |
| Linear track NEMA11 motor (50 mm) | Befenybay | BE073-1 | 47.3 | 3.00 | 141.90 |
| 3D-printed optical post holder | | | | 1.00 | 0.00 |
| 3D-printed breadboard to linear track | | | | 1.00 | 0.00 |
| **Speaker and Amplifier** | | | | | |

| | | | | | |
|---|---|---|---|---|---|
| Amplifiers | DROK | DROK | 10.75 | 1.00 | 10.75 |
| Speakers | Intervox | S150RMSS-R2W | 0.75 | 2.00 | 1.50 |
| L-Bracket | McMaster-Carr | 1556A54 | 1.06 | 1.00 | 1.06 |
| Optical Post | Thorlabs | TR3 | 6.58 | 2.00 | 13.16 |
| 3D-printed casing for speaker-amplifier | | | | 1.00 | 0.00 |
| **Stepper-motor drivers** | | | | | |
| Stepper-motor drivers | Banria | 4257 | 26.9 | 3.00 | 80.70 |
| Power supply | DROK | 24V | 15.89 | 1.00 | 15.89 |
| Stepper motor housing (laser cut) | | | | 1.00 | 0.00 |
| **Optogenetics-power modulation** | | | | | |
| Complete assembly | | | 25 | 1.00 | 25.00 |
| 3D printed housing | | | | 1.00 | 0.00 |
| **Miscellaneous** | | | | | |
| 4-40 Screws | Thorlabs | SH4S025 | 0.18 | 10.00 | 1.80 |
| 1/4" Screws | Thorlabs | SH25S025 | 0.55 | 10.00 | 5.50 |
| Table Clamp | Thorlabs | CL5 | 5.67 | 2.00 | 11.34 |
| BNC cables | 50 ohm | 3 feet | 5.63 | 5.00 | 28.15 |
| 22 AWG wiring | Tensility | 30-00689 | 2 | 10.00 | 20.00 |
| Banana plug | na | na | 1.7 | 1.00 | 1.70 |
| Computer | Dell | Optiplex 7060 | 276 | 1.00 | 276.00 |
| Screen | Dell | SE2422H 24" | 90 | 1.00 | 90.00 |
| 12" x 18" plate | | | | 1.00 | 0.00 |
| **Optional** | | | | | |
| Breadboard 12" x 18" x 1/2" | Thorlabs | MB1218 | 459.36 | 1.00 | 459.36 |
| Breadboard 6" round x 1/2" | Thorlabs MBR6 | MBR6 | 156.35 | 1.00 | 156.35 |
| **Total costs rig** | | | | | **1077.37** |
| **Including computer** | | | | | **1443.37** |
| **Including optional** | | | | | **2059.08** |

**Supplementary Figure 1. Itemized hardware cost estimate for the SPOUT platform.** Complete bill of materials for building one SPOUT rig, grouped by subsystem (breakout board, power supply, dual-lick detector, water reservoir/solenoids/spouts, head-restriction set-up, motorized manipulator, speaker and amplifier, stepper-motor drivers, optogenetic power modulation, and miscellaneous). For each item the supplier, model number, unit price, quantity, and line total are listed in US dollars. Parts fabricated in-house (3D-printed housings, laser-cut and CNC-cut components) are listed. The core rig totals approximately US $1,077, rising to US $1,443 with a dedicated computer and monitor, and to US $2,059 including the optional optical breadboards.

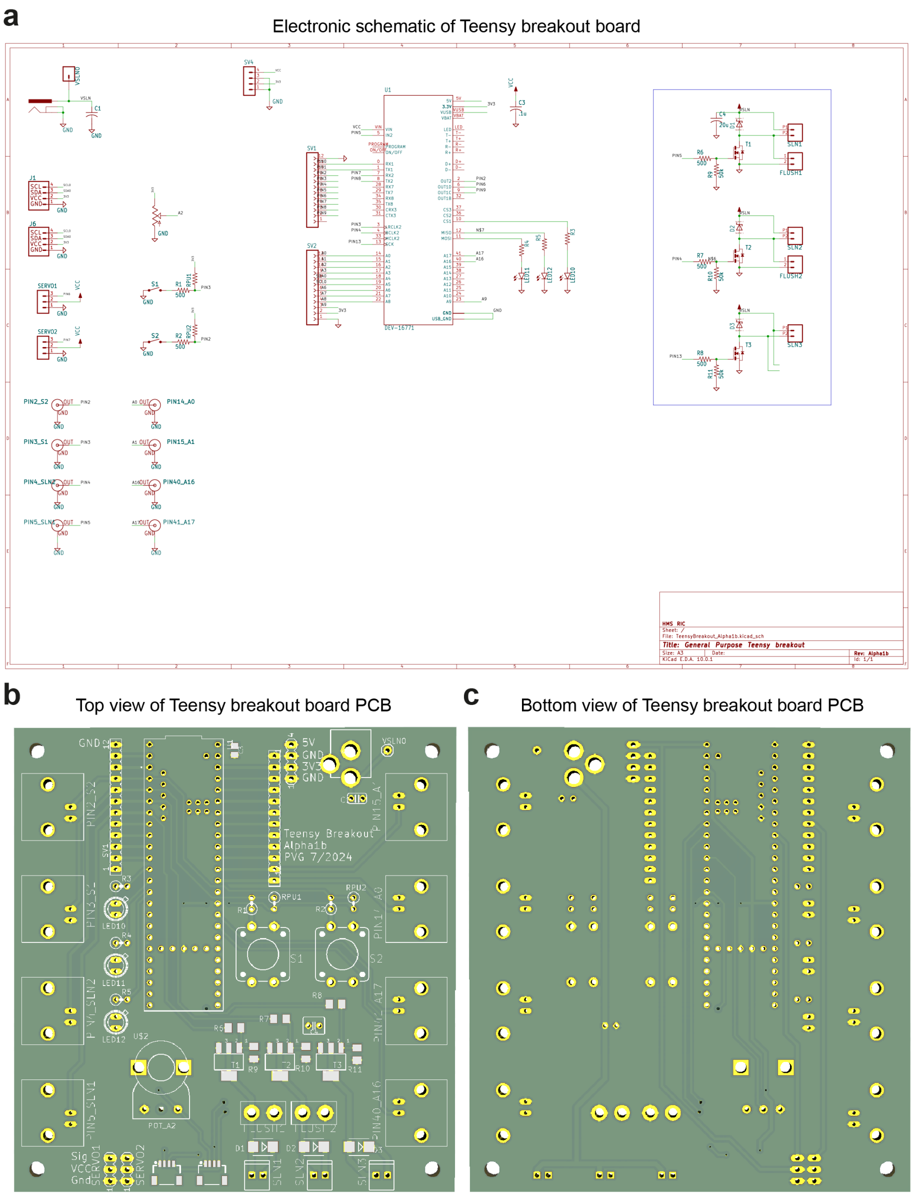
a
Electronic schematic of Teensy breakout board
Title: General Purpose Teensy breakout
b
Top view of Teensy breakout board PCB
Teensy Breakout
Alpha1b
PVG 7/2024
c
Bottom view of Teensy breakout board PCB

**Supplementary Figure 2. Design and layout of the in-house developed Teensy breakout board.** (A) Complete electronic schematic (KiCad) of the general-purpose Teensy breakout board, showing the Teensy microcontroller and its peripheral circuitry: the integrated solenoid drivers (blue outline), servo-motor connectors, potentiometer inputs, and the broken-out digital and analog input/output pins referenced throughout the platform. (B) Top-copper and (C) bottom-copper views of the fabricated printed circuit board (PCB), with labeled connector footprints and component placements.

**a**

Electronic schematic of dual-lick detector

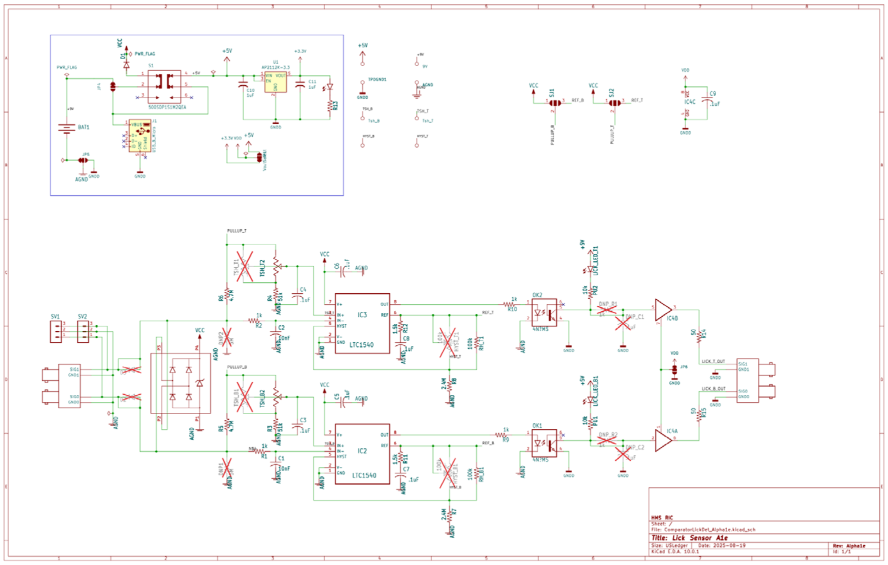


**b**

Top view of dual-lick detector PCB

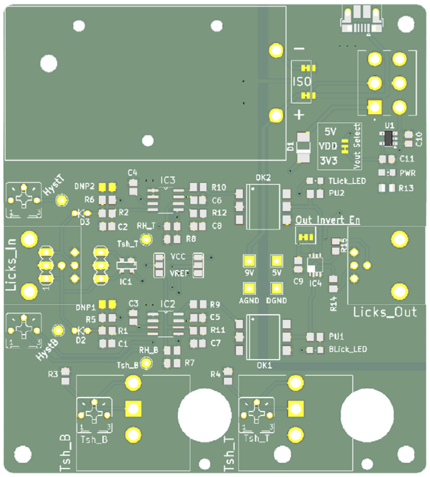


**c**

Bottom view of dual-lick detector PCB

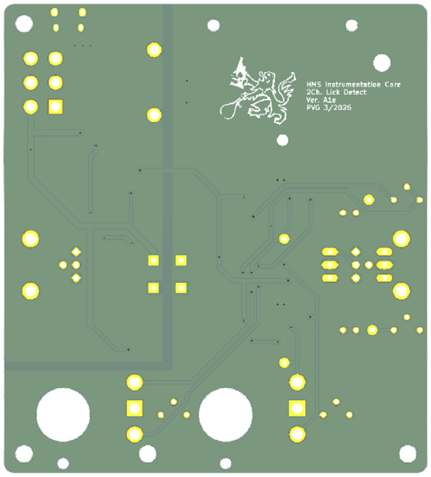

**Supplementary Figure 3. Design and layout of the conduction-based dual-lick detector.** (A) Complete electronic schematic (KiCad) of the two-channel lick detector, comprising two independent conduction-based sensing channels with adjustable threshold, selectable output inversion, and optical isolation of the sensing front end from the recording electronics. (B) Top-copper and (C) bottom-copper views of the fabricated PCB, with labeled lick inputs (Licks_In), buffered outputs (Licks_Out), threshold trimmers, and supply-voltage select jumpers.

**a**

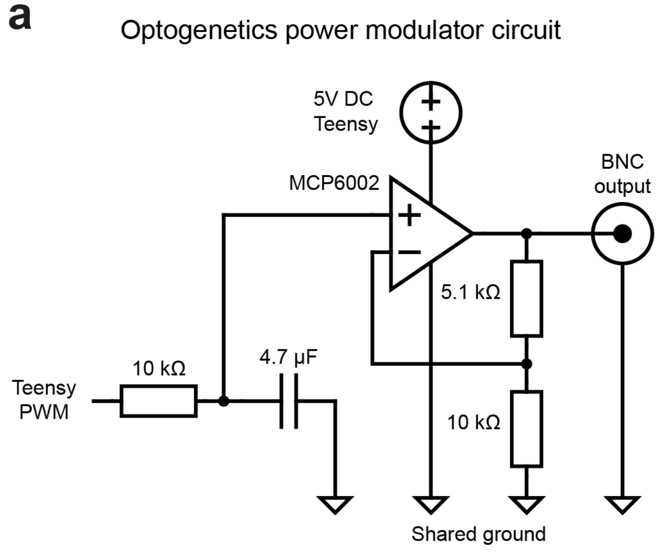
Optogenetics power modulator circuit
5V DC
Teensy
MCP6002
BNC
output
5.1 kΩ
10 kΩ
4.7 µF
Teensy
PWM
10 kΩ
Shared ground

**Supplementary Figure 4. Schematic of optogenetics power modulator circuit.** (A) Complete electronic schematic of the optogenetics power modulator that scales Teensy's native 3.3 V logic output to 5 V using a simple operational-amplifier (MCP6002). This circuit allows Teensy's pulse-width modulation to command a power.
